# SARS-CoV-2 receptor ACE2 and TMPRSS2 are predominantly expressed in a transient secretory cell type in subsegmental bronchial branches

**DOI:** 10.1101/2020.03.13.991455

**Authors:** Soeren Lukassen, Robert Lorenz Chua, Timo Trefzer, Nicolas C. Kahn, Marc A. Schneider, Thomas Muley, Hauke Winter, Michael Meister, Carmen Veith, Agnes W. Boots, Bianca P. Hennig, Michael Kreuter, Christian Conrad, Roland Eils

## Abstract

The SARS-CoV-2 pandemic affecting the human respiratory system severely challenges public health and urgently demands for increasing our understanding of COVID-19 pathogenesis, especially host factors facilitating virus infection and replication. SARS-CoV-2 was reported to enter cells via binding to ACE2, followed by its priming by TMPRSS2. Here, we investigate *ACE2* and *TMPRSS2* expression levels and their distribution across cell types in lung tissue (twelve donors, 39,778 cells) and in cells derived from subsegmental bronchial branches (four donors, 17,521 cells) by single nuclei and single cell RNA sequencing, respectively. While *TMPRSS2* is expressed in both tissues, in the subsegmental bronchial branches *ACE2* is predominantly expressed in a transient secretory cell type. Interestingly, these transiently differentiating cells show an enrichment for pathways related to RHO GTPase function and viral processes suggesting increased vulnerability for SARS-CoV-2 infection. Our data provide a rich resource for future investigations of COVID-19 infection and pathogenesis.

## INTRODUCTION

In December 2019, a disease affecting predominantly the respiratory system emerged in Wuhan, province Hubei, China, with its outbreak being linked to the Huanan seafood market as about 50% of the first reported cases either worked at or lived close to this market [Chen et al. (2020), Huang et al. (2020)]. Among the first 99 reported patients with an average age of 55.5 years, 2/3 were male and 50% suffered from chronic diseases [Chen et al. (2020)]. The disease rapidly spread to other provinces in China, neighboring countries and eventually worldwide [Wang et al. (2020a), Wu and McGoogan (2020), Zhu et al. (2020)]. The World Health Organization (WHO) named the disease **co**rona**vi**rus **d**isease 20**19**, COVID-19 (formerly known as 2019-nCov), and the virus causing the infection was designated as **s**evere **a**cute **r**espiratory **s**yndrome **co**rona**v**irus **2**, SARS-CoV-2 [Coronaviridae Study Group of the International Committee on Taxonomy of (2020)], belonging to the *Coronaviridae* family.

The two coronavirus infections affecting global public health in the 21^st^ century were caused by SARS-CoV and MERS-CoV (**M**iddle **E**ast **r**espiratory **s**yndrome **co**rona**v**irus) [de Wit et al. (2016)]. As suggested for SARS-CoV-2, these two coronaviruses had a zoonotic origin with human-to-human transmission after their outbreak [de Wit et al. (2016)]. SARS-CoV and MERS-CoV infected 8,096 and 2,494 humans with 10% and 37% deaths, respectively [WHO (2004), WHO (2016), de Wit et al. (2016)]. As of 13 March 2020, 128,392 SARS-CoV-2 infections were reported worldwide with 4,728 deaths. The mean incubation time is hard to be precisely defined during an ongoing outbreak, but currently predicted to rank form 4 - 6.4 days [Guan et al. (2020), Backer et al. (2020)] up to 14 days as estimated by the WHO for COVID-19 and thus very similar to those of SARS and MERS [Backer et al. (2020)]. Until 11 February, most of the COVID-19 patients showed only mild symptoms (80%) or none at all (1% confirmed cases) and 14% had severe symptoms, while 5% of the COVID-19 patients were critically affected [Wu and McGoogan (2020)]. These data suggest that the estimated number of undetected SARS-CoV-2 infections is much higher and even patients with mild or no symptoms are infectious [(Lai et al., 2020); Rothe et al. (2020)]. Deaths were solely reported for the critical cases (49%), mainly for patients with preexisting comorbidities such as cardiovascular disease, diabetes, chronic respiratory disease, hypertension or cancer [Wu and McGoogan (2020)]. Although COVID-19 has a milder clinical impairment compared to SARS and MERS for the vast majority of patients, SARS-CoV-2 infection shows dramatically increased human-to-human transmission rate with the total number of deaths significantly exceeding those of SARS and MERS patients already within the first three months of the COVID-19 outbreak. The emergent global spread of SARS-CoV-2 and its strong impact on public health immediately demands for joint efforts in bio-medical research increasing our understanding of the virus’ pathogenesis, its entry into the host’s cells and host factors facilitating its fast replication that explains the high human-to-human transmission rates.

SARS-CoV-2 is an enveloped virion containing one positive-strand RNA genome and its sequence has already been reported [Chan et al. (2020), Lu et al. (2020), Wu et al. (2020a), Wu et al. (2020c), Zhou et al. (2020)]. The genome of SARS-CoV-2 comprises 29.9 kb and shares 79.5% and 96% identity with SARS-CoV and bat coronavirus, respectively [Zhou et al. (2020)]. Coronaviruses were reported to use different cellular entry mechanisms in terms of membrane fusion activities after receptor binding [White and Whittaker (2016)]. SARS-CoV was previously shown to bind to angiotensin-converting enzyme 2 (ACE2) for cell entry, mediated via the viral surface spike glycoprotein (S protein) [Gallagher and Buchmeier (2001), Li et al. (2003), Simmons et al. (2013)]. Comparison of the SARS-CoV and SARS-CoV-2 S protein sequence revealed 76% protein identity [Wu et al. (2020b)] and recent studies reported that SARS-CoV-2 is also binding to ACE2 *in vitro* [Hoffmann et al. (2020), Walls et al. (2020), Yan et al. (2020), Zhou et al. (2020)]. Subsequently, the S protein is cleaved by the transmembrane protease serine 2 TMPRSS2 [Hoffmann et al. (2020)]. Simultaneously blocking TMPRSS2 and the cysteine proteases CATHEPSIN B/L activity inhibits entry of SARS-CoV-2 *in vitro* [Kawase et al. (2012)], while the SARS-CoV-2 entry was not completely prohibited *in vitro* [Hoffmann et al. (2020)]. It remains to be determined whether additional proteases are involved in priming of the SARS-Cov-2 but not SARS-CoV S protein. One candidate is FURIN as the SARS-CoV-2 S protein contains four redundant FURIN cut sites (PRRA motif) that are absent in SARS-CoV. Indeed, prediction studies suggest efficient cleavage of the SARS-CoV-2 but not SARS-CoV S protein by FURIN [Coutard et al. (2020), Wu et al. (2020b)]. Additionally, FURIN was shown to facilitate virus entry into the cell after receptor binding for several coronaviruses, e.g. MERS-CoV [Burkard et al. (2014)] but not SARS-CoV [Millet and Whittaker (2014)]. In addition, although *ACE2* was previously described to be expressed in the respiratory tract [Jia et al. (2005), Ren et al. (2006), Xu et al. (2020), Zhao et al. (2020), Zou et al. (2020)], still little is known about the exact cell types expressing *ACE2* and *TMPRSS2* serving as entry point for SARS-CoV and the currently emerging SARS-CoV-2. Therefore, there is an urgent need for investigations of tissues in the upper and lower airways in COVID-19 patients but also healthy individuals to increase our understanding of the host factors facilitating the virus entry and its replication, ultimately leading to treatment strategies of SARS-CoV-2 infections.

As pointed out recently [Zhang et al. (2020b)], our understanding of host genetic factors involved in COVID-19 disease outcome is still poor.

## RESULTS

### *ACE2* and its co-factor *TMPRSS2* are expressed in the lung and bronchial branches

Here, we established a rich reference data set that describes the transcriptional landscape at the single cell level of the lung and subsegmental bronchial branches of in total 16 individuals (Fig. 1A). Based on this resource, we set out to identify potential key mechanisms likely involved in the SARS-CoV-2 pathway. First, we investigated the expression patterns of the SARS-CoV-2 receptor *ACE2* and the serine protease priming its S-protein, *TMPRSS2*, in individual cells in the lung and in subsegmental bronchial branches (Fig. 1A).

**Figure 1.**
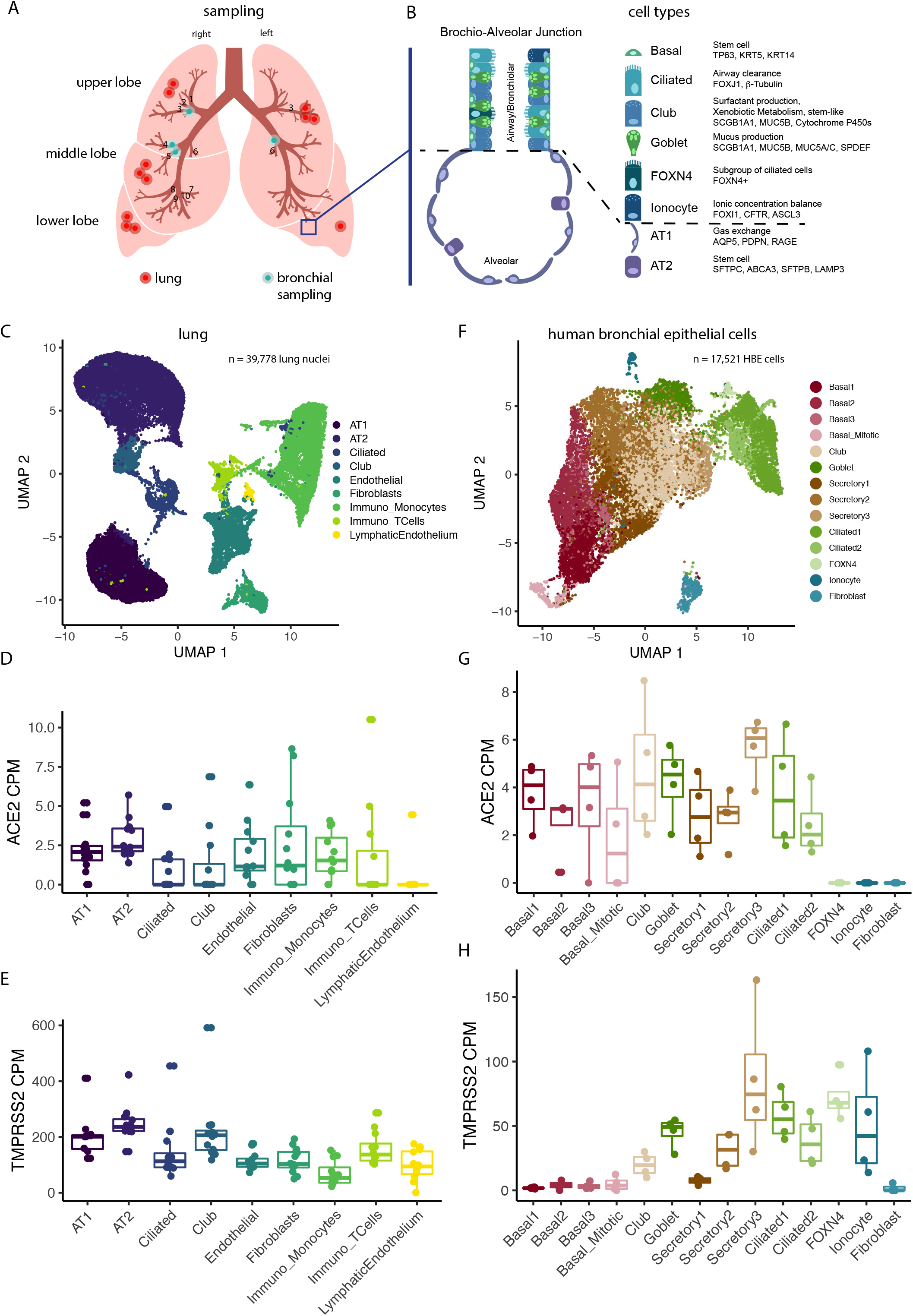
*ACE2* and *TMPRSS2* are expressed in specific cell types in lungs and HBECs. A) Sampling location of the surgical lung specimens and human bronchial epithelial cells (HBECs) used in this study. Blue rectangle is zoomed in B) Overview of the major cell types in the lung and airways. C) Uniform manifold approximation and projection (UMAP) of primary lung samples single nuclei RNA sequencing (cell types are color-coded). D) Expression values of *ACE2* in the cell types of primary lung samples. E) Expression values of *TMPRSS2* in the cell types of primary lung samples. F) UMAP projections of HBE single cell RNA sequencing data (cell types are color-coded), G) Expression values of *ACE2* in the cell types of HBECs. H) Expression values of *TMPRSS2* in the cell types of HBECs. Boxes in box plots indicate the first and third quartile, with the median shown as horizontal lines. Whiskers extend to 1.5 times the inter-quartile-range. All individual data points are indicated on the plot.

To quantify gene expression in the lung, single nuclei RNA sequencing was performed on surgical specimens of healthy, non-affected lung tissue from twelve lung adenocarcinoma (LADC) patients, resulting in 39,778 sequenced cell nuclei. All major cell types known to occur in the lung were identified (Fig. 1B, C). Independent of the cell types present in the lung, the median *ACE2* levels were below five counts per million (CPM) (Fig. 1D), which given a typical mRNA content of 500,000 mRNA molecules per cell indicate that only about half of all cells were statistically expected to contain even a single *ACE2* transcript. The reads per patient and cell type were therefore aggregated into pseudo-bulks and analysis was continued. As expected from prior literature, the AT2 cells showed highest *ACE2* expression in the lung both in terms of their CPM values (Fig. 1D, further referred to as *ACE2*^+^ cells) and the percentage of positive cells (Supp. Fig. 3). The expression of *TMPRSS2* across cell types of the lung (further referred to as *TMPRSS2*^+^ cells) was much stronger with a certain specificity for AT2 cells (Fig. 1E), which is in agreement with previous studies.

For the subsegmental bronchial branches, air-liquid-interface (ALI) cultures were grown from primary human bronchial epithelial cells (HBECs) and subjected to single cell RNA sequencing, resulting in 17,451 cells across four healthy donors (further referred to as HBECs). In this dataset, we identified all expected cell types, including recently described and rare cell types such as *FOXN4*-positive cells and ionocytes (Fig. 1F, Supp. Fig. 1, 2). Expression levels of *ACE2* were comparable but slightly elevated compared to the lung tissue samples, with the strongest expression being observed in a subset of secretory cells (‘secretory3’; Fig. 1G). While *TMPRSS2* was less strongly expressed in HBECs than in lung tissue cells, its expression was still markedly higher than that of *ACE2* (Fig. 1H). Interestingly, *ACE2*^+^ cells were also most strongly expressing *TMPRSS2* (Fig. 1F, H). In the HBECs, a significant enrichment of the number of *ACE2*^+^/*TMPRSS2*^+^ double-positive cells was observed (1.6 fold enrichment, p=0.002, Supp. Fig. 4).

### Transient secretory cells display a transient cell state between goblet and ciliated cells

As the ‘secretory3’ cells in the HBECs showed comparably strong expression of both genes implicated in SARS-CoV-2 entry, with *ACE2* expression levels in a similar to higher range than those in AT2 cells, we decided to further investigate this cell type. As airway epithelial cells are known to continuously renew in a series of differentiation events involving nearly all major HBE cell types in a single trajectory, we mapped the ‘secretory3’ cells according to their position in this hierarchy using pseudo-time mapping (Fig. 2A). ‘Secretory3’ cells were found at an intermediate position between club or goblet cells and ciliated cells (Fig. 2A), which could be confirmed by shared marker gene expression (Fig. 2B). This placed ‘secretory3’ cells at a point shortly before terminal differentiation is reached (Fig. 2C, D). Hence, we will refer to these cells as ‘transient secretory cells’.

**Figure 2.**
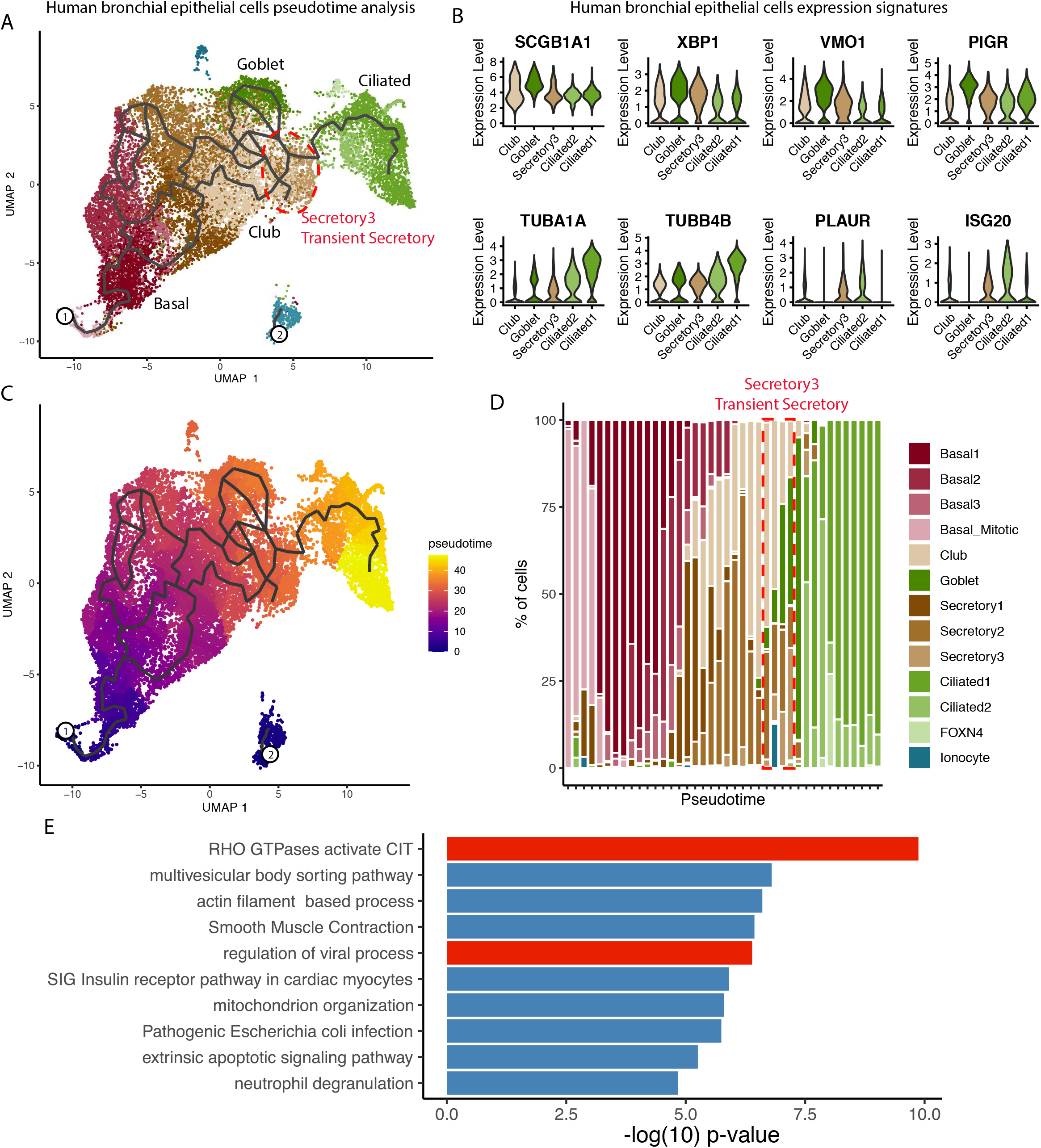
Characterization of transient secretory (secretory3) cells. A) Pseudotime trajectory projected onto a UMAP embedding of HBECs. The location of the secretory3 cell type is marked by a red outline (cell types are color-coded). B) Expression of goblet (top row) and ciliated cell markers (bottom row) in club, goblet, secretory3, and ciliated cells. C) Pseudo-time trajectory projected onto a UMAP embedding of HBECs (pseudo-time values are color-coded). D) Cell type composition along a binned pseudo-time axis. The earliest stages are located on the left. E) Pathway enrichment values for secretory3-specific marker genes.

In agreement with ongoing differentiation towards an epithelial cell lineage, the cellspecific markers of the transient secretory cells showed an enrichment of RHO GTPases and their related pathways (Fig. 2E). Note that we find this enrichment of pathways exclusively in the transient secretory cells but in none of the other cell types. As RHO GTPases have been implicated in membrane remodeling and the viral replication cycle, especially entry, replication, and spread [Van den Broeke et al. (2014)], transient secretory cells may be more permissive to SARS-CoV-2 infection adding to a potential vulnerability caused by considerably high co-expression levels of *ACE2* and *TMPRSS2*. Interestingly, these cell marker genes were not enriched in any of the cell types in the whole lung indicating the absence of this transient secretory cell type in the lung samples. This may indicate a different entry route for the SARS-CoV-2 virus in the more central bronchial branches compared to the peripheral lung tissue.

### FURIN expressing cells overlap with *ACE2*^+^/*TMPRSS2*^+^ and *ACE2*^+^/*TMPRSS2*^-^ cells in lung and in bronchial cells

We next sought to explore whether we could derive additional evidence for other factors recently suggested to be involved in SARS-CoV-2 host cell entry. Therefore, we investigated the expression of *FURIN*, as SARS-CoV-2 but not SARS-CoV was reported to have a FURIN cleavage site in its S protein, potentially increasing its priming upon ACE2 receptor binding. In lung tissue, AT2, endothelial, and immune cells were most strongly positive for *FURIN* expression (Fig. 3A). Across cell types, we observed a marked enrichment of the number of double and triple positive cells for any combination of *ACE2*, *TMPRSS2*, and/or *FURIN* expression, indicating a preference for co-expression (Fig. 3B). Interestingly, also transient secretory cells in the HBECs showed an intermediate expression of *FURIN* (Fig. 3C). While *TMPRSS2* and *FURIN* were less strongly expressed in HBECs than in distant cells from surgical lung tissue, their expression was still markedly higher compared to *ACE2*. In the HBECs, a significant enrichment of the number of *ACE2*^+^/*TMPRSS2*^+^ double-positive cells was observed, while *ACE2*^+^/*TMPRSS2*^+^/*FURIN*^+^ enrichment did not reach significance (Fig. 3D). The latter finding, however, may be due to differences in coexpression or the generally lower number of positive cells reducing statistical power. Interestingly, our data showed that the additional possibility of priming the SARS-CoV-2 S protein by FURIN would potentially render 25% more cells vulnerable for infection as compared to by exclusively TMPRSS2 S protein priming. Thus, our data suggest that FURIN might increase overall permissiveness of cells in the respiratory tract by potentially equipping more cells with proteolytic activity for SARS-CoV-2 S protein priming after ACE2 receptor binding.

**Figure 3:**
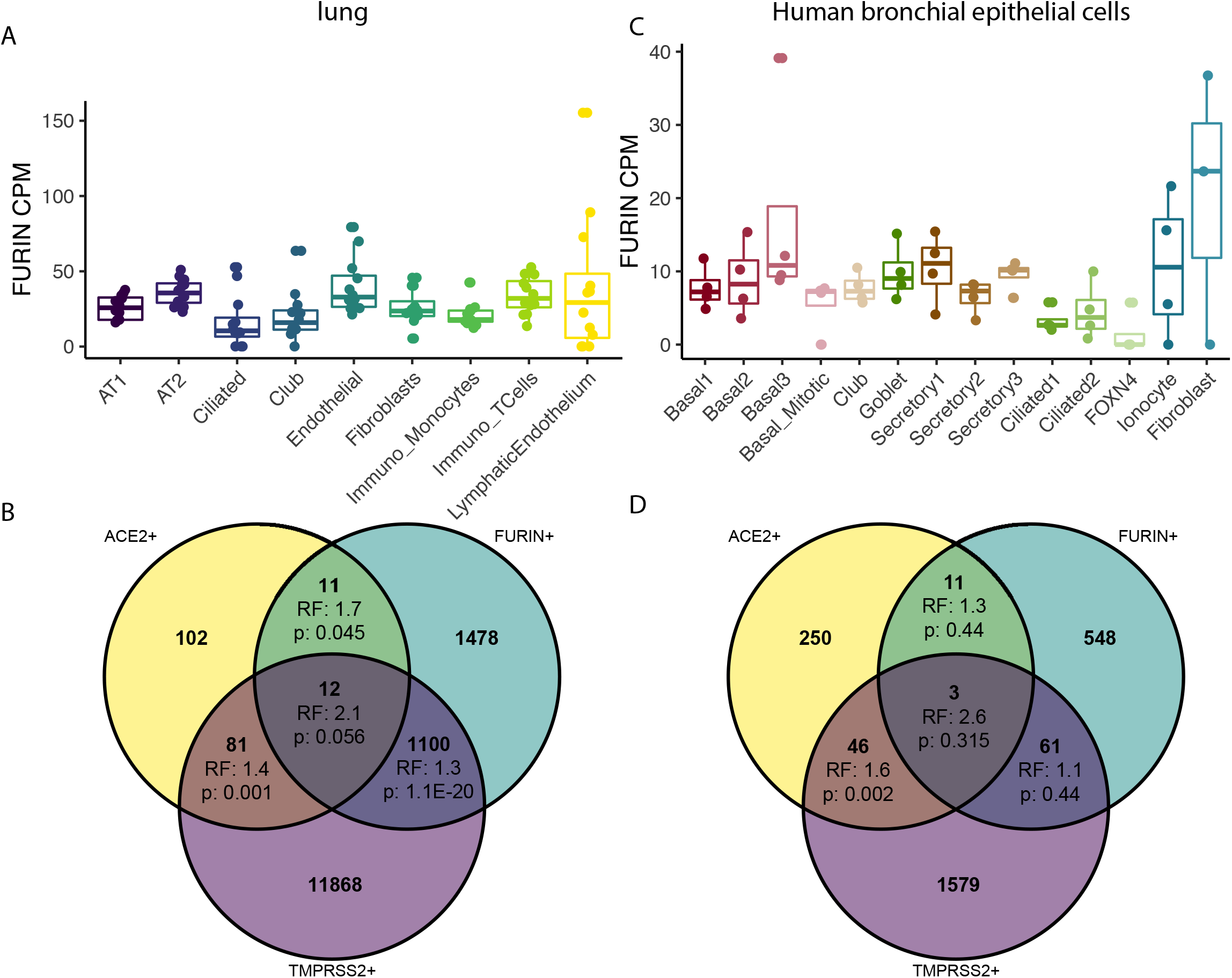
*FURIN* is expressed in *ACE2*^+^ and *ACE2*^+^/*TMPRSS2*^+^ cells. A) Expression values of *FURIN* in the cell types of primary lung. B) Overlap of *ACE2*^+^, *TMPRSS2*^+^, and *FURIN*^+^ cells in the lung dataset. RF: representation factor, enrichment. P: hypergeometric tail probability. Total number of cells: 39,778. C) Expression values of *FURIN* in HBECs. D) Overlap of *ACE2*^+^, *TMPRSS2*^+^, and *FURIN*^+^ cells in the HBEC dataset. RF: representation factor, enrichment. P: hypergeometric tail probability. Total number of cells: 17,451. Boxes in box plots indicate the first and third quartile, with the median shown as horizontal lines. Whiskers extend to 1.5 times the inter-quartile-range. All individual data points are indicated on the plot.

### Sex, age and history of smoking in correlation with *ACE2* expression

Early epidemiological data on SARS-CoV-2 transmission and spread have suggested age, sex and history of smoking among others as potential confounding factors impacting SARS-CoV-2 infection [Brussow (2020), Chen et al. (2020), Huang et al. (2020), Zhang et al. (2020a)]. We thus investigated these possible risk factors for COVID-19 and their influence on receptor gene expression (Fig. 4A). No sex-related difference could be observed in the *ACE2* expression in individual cell types neither in the lung tissue nor in the HBECs (Fig. 4C, D). This observation is in full agreement with recent reports based on a large number of COVID-19 patients [Guan et al. (2020)]. Although the first reported cases suggested higher infection rates for males [Chen et al. (2020), Huang et al. (2020)], no significant differences in the infection rate of males and females were found with increasing numbers of COVID-19 patients [Wang et al. (2020b), Brussow (2020), Zhang et al. (2020a)].

**Figure 4.**
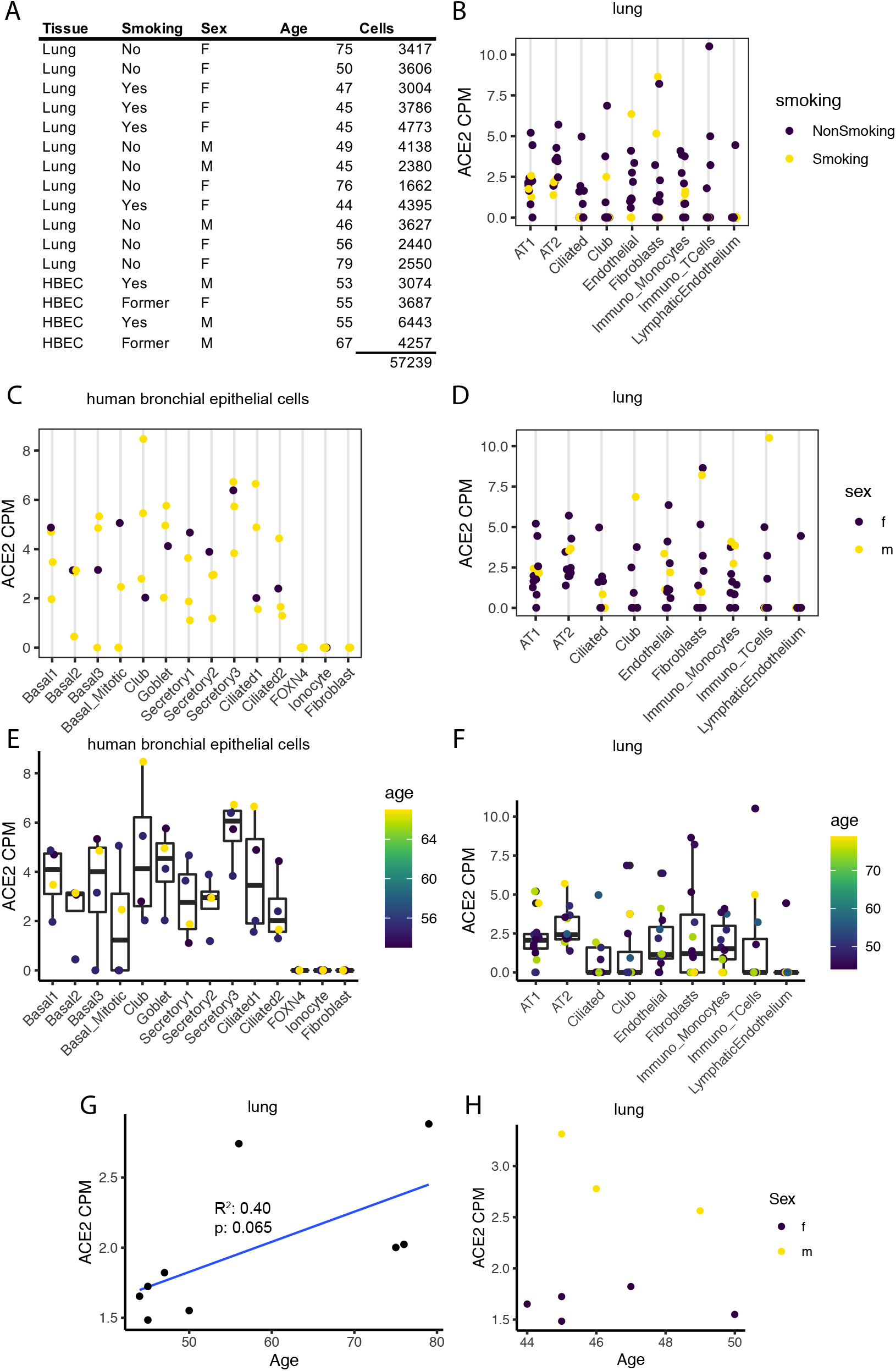
Age, sex and smoking behavior as possible factors influencing *ACE2* expression. A) Metadata of the sample donors. B) Expression of *ACE2* in the cell types of the primary lung (history of smoking color-coded). C) and D) Expression of *ACE2* in C) HBECs and D) primary lung (sex is color-coded). E) and F) Expression of *ACE2* in E) HBECs and F) primary lung (age is color-coded). G) and H) *ACE2* expression over all cell types versus age in G) female primary lung cells and H) primary lung cells of patients aged 40 to 50 (sex is color-coded). Boxes in box plots indicate the first and third quartile, with the median shown as horizontal lines. Whiskers extend to 1.5 times the inter-quartile-range. All individual data points are indicated on the plot.

It has to be noted, though, that the power of our datasets to detect such changes with lung cells from surgical lung tissue samples from twelve patients (three male, nine female) and HBECs from four patients (three male, one female) is limited. On the individual cell type level, no *ACE2* expression differences correlating with age could be observed (Fig. 4E, F), and neither did the cell type composition change notably (Supp. Fig. 6). Although no dependency of *ACE2* expression on age, gender and sex was found in single cell populations, we observed interesting trends in the lung for age and gender dependencies when aggregating all reads per sample into a single *ACE2* expression value (Figure 4G, H). Here, we see a trend for increasing overall expression of *ACE2* with age for all female lung samples (R^2^=0.40; p=0.065). Since we only have samples from males of younger age available, we cannot study age dependencies for males here. However, when comparing *ACE2* expression of all five female and three male samples in the young age group between 40 and 50 years of age, we see a clearly higher overall expression of *ACE2* in males compared to females (p=0.048). Note that smoking did not seem to affect *ACE2* expression levels in the lung cells within our dataset as all smokers in our study were of about the same age and *ACE2* expression correlated with age (Fig. 4B).

Altogether, we here demonstrate how single cell sequencing studies of samples from anatomically precisely defined parts of the respiratory system can provide new insights into potential host cell vulnerability for SARS-CoV-2 infection.

## DISCUSSION

The here presented transcriptome data on single cell level of healthy human lung tissues, including surgical lung specimen and subsegmental bronchial branches, contains valuable information about human host factors for SARS-CoV-2 infection and other related infectious diseases affecting the respiratory tract. This resource certainly comes with limitations (see below). However, we believe the unprecedented depth of our analysis on a single cell level will provide a valuable resource for future mechanistic studies and target mining for pulmonary host factors that are involved in facilitating virus entry and replication, ultimately leading to defining genes that are of urgent interest for studying transcriptional changes in COVID-19 patients or the SARS-CoV-2 pathogenicity in general.

This data set comprises 16 individuals and a total of 57,229 cells. Thus, this large single-cell and nuclei expression resource enables the analysis of weakly expressed genes such as *ACE2* and identification of rare cell types and the transient secretory cell type, for which our data showed a particularly high level of potential vulnerability for SARS-CoV-2 infection assuming that ACE2 is the receptor that is likely to be crucial for its cell entry. Although we strongly believe that the here presented data is well suited as reference data set to study SARS-CoV-2 infection on the single cell transcriptional level, we are also well aware of potential limitations of our data.

First, the here presented data is derived from individuals that have no infection history with SARS-CoV-2. However, we deem our data clinically meaningful as our patient cohort is representative for those being affected by SARS-CoV-2 associated disease. The patient cohort analyzed here is in line with recently published data on characteristics of COVID-19 patients with regards to age (this study: median age 48 years - lung cohort or 55 years - HBEC cohort; Chen *et al.,* 2020: 55 years; Guan *et al.,* 2020: 47 years) [Chen et al. (2020), Guan et al. (2020)] (Guan NEJM, Chen Lancet) and especially comorbidity burden [Chen et al. (2020)] with 50% of the patients being affected by chronic cardiovascular or pulmonary disease or diabetes (this study: 50%; Chen *et al.,* 2020: 51%; Guan *et al.,* 2020: no data available). However, there was a different prevalence of smoking or history of smoking in our cohort (this study: 25% - lung cohort or 100% HBEC cohort; Guan *et al.,* 2020: 85%)

Second, we are combining data of different sections of the lung and use two different RNA sequencing methods on single cell level. While single cell RNA sequencing typically delivers higher quality data, the cells must be intact for processing, as damaged cells present a loss of RNA molecules with mitochondrial reads preferentially retained, introducing a skewed expression profile [Luecken and Theis (2019)]. Adequate sampling of lung specimen for preparing suspensions of intact cells is often challenging. Single nuclei RNA sequencing also reveals transcriptome data on single cell level and can be used on frozen tissue. This method typically provides a more faithful estimate of cell type proportions due to lessened dissociation bias. However, it typically comes with less reads per cell and less genes detected [Bakken et al. (2018)], although this bias seems low in our dataset (Supp. Fig. 7). The identification of closely related cell populations is mostly concordant in both methods, with single cell RNA sequencing being inferior for detection of lowly expressed genes on a single-cell level [Bakken et al. (2018)]. The gene expression shown here for cells from surgical lung tissue samples and ALI-cultured cells derived from the subsegmental bronchial tree (HBECs) should not be directly compared, as the different techniques are likely to have an influence on absolute expression values. However, qualitative comparisons of relative abundances as performed here are still appropriate.

Third, the number of samples included here is very much limiting the scope of understanding the pathogenicity of SARS-CoV-2 in the context of different confounding factors such as age, gender and history of smoking. As a consequence of small sample numbers, the significance level for dependency of expression of single genes, most predominantly *ACE2*, on such confounding factors are at best weak. However, we believe that it is a strength of our data that our clinical samples are well annotated w.r.t. anatomical position and patient characteristics. Thus, our data is in principle suited for testing hypothesis derived from larger epidemiological studies. Clearly, measuring expression levels for specific genes in larger patient cohorts is required to further substantiate the hypothesis derived from our and others’ data.

Bearing the above limitations in mind, we investigated *ACE2* expression levels in all cell types in the lung and bronchial branches revealing very low expression levels that are nevertheless significantly enriched in AT2 cells in the lung as described previously [Hamming et al. (2004)] and are comparable with the levels in specific cell types in the bronchial branches. As ACE2 is the only currently known SARS-CoV-2 receptor on the host cell surface, we propose that higher expression levels of *ACE2* facilitate infection by SARS-CoV-2.

While we did not find any dependency of *ACE2* expression on sex, gender and age on the single cell level, we observed a trend for age dependency on *ACE2* expression aggregated over all cell types in lung samples from females. Recent data suggest that infection rates are similar in children as in adults, but remains undiagnosed since the clinical picture is often subclinical [Bi et al. (2020)]. Hence, it would be of utmost interest to study both healthy and infected children to understand the transcriptional basis accounting for differences in development of clinical symptoms. Furthermore, analyzing further samples from both males and females, both healthy and infected, of different age groups on the single cell level will lay the foundation for understanding vulnerability of different cell types for SARS-CoV-2 infection in different parts of the respiratory system.

One emergent question is why the human-to-human transmission of SARS-CoV-2 is much higher compared to SARS-CoV or MERS-CoV. Potential explanations comprise i) the binding of SARS-CoV-2 to another, yet unknown receptor on the host cell surface, ii) enhanced cleavage of the SARS-CoV-2 S protein resulting in higher efficiency of the virus’ entry into the cell, and iii) additional host factors increasing the virus entry into the cell, e.g. by facilitating membrane fusion. We used our reference map and our above described findings, seeking for additional host factors that might be involved in any of those mechanisms.

Coronaviruses were shown to be able to enter into the host cell via several pathways, including endosomal and non-endosomal entry in the presence of proteases [Simmons et al. (2004), Matsuyama et al. (2005), Wang et al. (2008), White and Whittaker (2016)]. Different proteases were previously shown to facilitate entry into the cells for different coronaviruses [Hamming et al. (2004)]. Both SARS-CoV and SARS-CoV-2 were shown to share the ACE2 as cell surface receptor and TMPRSS2 as the major protease facilitating their entry into the host cell [Hoffmann et al. (2020), Walls et al. (2020), Yan et al. (2020), Zhou et al. (2020)]. Nevertheless, SARS-CoV-2 human-to-human transmission rates appear to be significantly higher as compared to of SARS-CoV, given the massively higher transmission rate three months after its first detection, with about 130,000 confirmed infections and more than 4,700 deaths (as compared to 8,096 SARS patients with 2,494 deaths from 2002-2003).

SARS-CoV-2 was shown to comprise a FURIN cleavage site that is absent in SARS-CoV [Coutard et al. (2020), Wu et al. (2020b)]. In addition to the presence of this FURIN cleavage site and experimental evidence that MERS-CoV is primed by FURIN, also recent data published by Hoffmann *et al.* (2020) suggest an additional protease besides TMPRSS2 and CATHEPSIN B/L as blockage of both completely prohibited the entry of SARS-CoV but not SARS-CoV-2. However, the involvement of FURIN in the virus infection pathway remains to be confirmed by future experimental approaches. Further evidence for an involvement of FURIN in the entry of SARS-CoV-2 into the host cell comes from very recent studies predicting increased binding affinity between the SARS-CoV-2 S protein and the human ACE2 receptor as compared to the SARS-CoV S protein [Wrapp et al. (2020), Wu et al. (2020b)]. Wu *et al.* postulate that the cleavage of the S protein at the FURIN cut site might cause the increased binding affinity of SARS-CoV-2 to its receptor, e.g. triggered by structural rearrangements of the cleaved S protein as it was previously shown for other coronavirus S proteins [Kirchdoerfer et al. (2016), Walls et al. (2016), Wrapp et al. (2020)].

As the involvement of an additional protease during S protein priming might enhance entry of SARS-CoV-2 compared to SARS-CoV, we also investigated co-expression of *ACE2*, *TMPRSS2*, and *FURIN*. Using the here presented reference data set, we were able to show co-expression of *ACE2* with *TMPRSS2* and/or *FURIN* in both healthy human lung tissue and HBECs. The potential additional priming of 25% of cells for SARS-CoV-2 infection by FURIN may support the hypothesis for increased human-to-human transmission rates. This hypothesis, however, must be followed up by future studies.

Besides higher SARS-CoV-2 infection rate of the cells due to FURIN cleavage, also the severity of COVID-19 in some of the clinical cases might be explained by this additional FURIN motif. Recent prediction studies suggest efficient cleavage of the SARS-CoV-2 S protein by FURIN that might result in higher pathogenicity of the virus [Coutard et al. (2020), Wu et al. (2020b)], potentially due to an increased affinity to the ACE2 receptor [Wrapp et al. (2020), Wu et al. (2020b)]. Further evidence for correlation of increased virus pathogenicity with FURIN priming arises from the observation that the presence of a FURIN-like cleavage site positively correlated with pathogenicity for different influenza strains [Kido et al. (2012)].

MERS-CoV is an additional coronavirus that comprises a FURIN-like motif [Burkard et al. (2014)]. Although this virus did not infect as many individuals as SARS-CoV or the currently emerging SARS-CoV-2, it was highly lethal (37% of MERS patients) in 2016 [WHO (2016), de Wit et al. (2016)]. However, although both SARS-CoV-2 and MERS-CoV are likely to be cleaved by FURIN during S protein priming, we speculate that the MERS-CoV receptor DPP4 does not have any or high relevance for COVID-19 as DPP4 is solely expressed in lung cells but not in the subsegmental bronchial branches (Supp. Fig. 5). Taken together, these data offer the interesting hypothesis that both high human-to-human transmission rate of SARS-CoV-2 and the severe COVID-19 cases might be caused by additional cleavage sites, resulting in higher ACE2 binding affinity and/or more efficient membrane fusion.

The most striking finding of this study was the detection of a specific cell type in the bronchial branches expressing *ACE2,* i.e. the transient secretory cells. Subsequent cell type marker analysis revealed that these cells reflect differentiating cells on the transition from club or goblet to terminally differentiated ciliated cells, the latter having an important role in facilitating the clearance of viral particles [Dumm et al. (2019), Sims et al. (2008)]. RHO GTPase activating CIT and viral processes regulating pathways are strongly enriched in the transient secretory cells. Interestingly, about 40% of the *ACE2^+^* transient secretory cells are co-expressing the protease *TMPRSS2* that is known to be involved in S protein priming, making this cell type potentially highly vulnerable for SARS-CoV-2 infection. If we include FURIN as another protease priming in SARS-CoV-2 entry of the host cell, the percentage of transient secretory cells expressing the receptor *ACE2* and either both or one of the proteases *TMPRSS2* and *FURIN,* increases to 50%. Hence, transient secretory cells would be potentially highly vulnerable for SARS-CoV-2 infection.

Interestingly, SARS-CoV replication was reported to be 100- to 1,000-fold higher via the non-endosomal pathway as compared to the endosomal pathway, with the non-endosomal pathway being dependent on the presence of proteases [Matsuyama et al. (2005)]. Therefore, our data suggest to experimentally investigate whether SARS-CoV-2 is able to enter into the cell also via the non-endosomal pathway, either by the involvement of several proteases or by the ongoing membrane remodeling in the transient secretory cells, which might include RHO GTPases as one prominent pathway specifically enriched in this cell type.

Taken together, we present a rich resource for studying transcriptional regulation of SARS-CoV-2 infection, which will serve as reference dataset for future studies of primary samples of COVID-19 patients and *in vitro* studies addressing the viral replication cycle. With all limitations in mind, which we discussed above, we demonstrate the potential of this resource for deriving novel and testing existing hypotheses that need to be followed up by independent studies.

## ACKNOWLEDGEMENTS

Cryopreserved surgical lung tissues from patients were kindly provided from the Lung Biobank Heidelberg, a member of the accredited Tissue Bank of the National Center for Tumor Diseases (NCT) Heidelberg, the BioMaterialBank Heidelberg and the Biobank platform of the German Center for Lung Research (DZL). We thank Martin Fallenbüchel and Christa Stolp for collection of tissue samples and Elizabeth Chang Xu for establishment of ALI cultures. We thank Leif Erik Sander, Irina Lehmann, and Saskia Trump for advice in this study. This study was supported by the European Commission (ESPACE, 874710, Horizon 2020) and the German Center for Lung Research (DZL, 82DZL00402). This publication is part of the Human Cell Atlas – www.humancellatlas.org/publications.

## AUTHOR CONTRIBUTIONS

N.C.K, M.A.S., M.M., C.C., and R.E. conceived and designed the project. N.C.K, M.A.S., M.M., A.W.B., B.P.H., M.K., C.C., and R.E. supervised the project. R.L.C, T.T., M.A.S., and C.V. performed experiments. S.L., R.L.C., T.T., and B.P.H. analyzed data. N.C.K., T.M., H.W., C.V., A.W.B. provided tissue and cells. S.K., M.A.S, T. M., B.P.H., M.K. C.C., and R.E. wrote the manuscript, all authors read and revised the manuscript.

## DECLARATION OF INTERESTS

The authors declare no competing interests.

## METHODS

### LEAD CONTACT AND MATERIALS AVAILABILITY

Further information and requests for resources and reagents should be directed to and will be fulfilled by the lead contact Roland Eils.

Primary lung and subsegmental bronchial samples are available upon request and upon installment of a material transfer agreement (MTA).

There are restrictions to the availability of the dataset due to potential risk of deidentification of pseudonymized RNA sequencing data. Hence, the raw data will be available under controlled access. Accession numbers, DOIs, or unique identifiers for raw data will be listed here upon submission to the community-endorsed public repository.

Processed data in the form of count tables and metadata tables containing patient ID, sex, age, smoking status, cell type and QC metrics for each cell are available on FigShare (https://doi.org/10.6084/m9.figshare.11981034.v1) and Mendeley Data (https://data.mendeley.com/datasets/7r2cwbw44m/1).

### EXPERIMENTAL MODEL AND SUBJECT DETAILS

#### Human lung tissues and bronchial branches

All subjects gave their informed consent for inclusion before they participated in the study. The study was conducted in accordance with the Declaration of Helsinki. The use of biomaterial and data for this study was approved by the local ethics committee of the Medical Faculty Heidelberg (S-270/2001 and S-538/2012).

Cryoconserved surgical healthy lung tissue from twelve patients was provided by the Lung Biobank Heidelberg.

Sex, age and smoking behavior for every individual is provided in figure 4a and additional information can be found in table 1.

**Table 1.** Patient characteristics. ELF: Epithelial lining fluid; ECOG: Eastern Cooperative Oncology Group Performance Status. Scale Data are expressed as range (mean ± SD). (* P < 0.05, *** P < 0.001).

|  | **Surgically lung patients** | **ELF sampled patients** |
| --- | --- | --- |
| **Number and gender (male/female)** | 12 (3/9) | 4 (3 / 1) |
| **Age (years)** | 44-79 (48 ± 13) | 53-67 (55 ± 6.4) |
| **ECOG** |  |  |
| 0 | 12 | 3 |
| 1 | 0 | 1 |
| **Smoking status** |  |  |
| *current smokers* | 3 | 2 |
| *ex-smokers* | 0 | 2 |
| *never smokers* | 9 | 0 |

### METHOD DETAILS

#### Lung tissues

Tissue was assembled during routine surgical intervention from lung cancer patients. A representative part of normal lung tissue distant from the tumor was cut into pieces of about 0.5-1 cm^3^ immediately after resection. Up to 4 pieces were distributed in 2 ml cyro vials (Greiner Bio-One GmbH). Thereafter, the vials were snap frozen in liquid nitrogen (within 0.5 h after resection). The tissue samples had no direct contact to the liquid liquid nitrogen. After snap-freezing, the vials were stored at −80°C until use [Muley et al. (2012)]. Patients’ characteristics are shown in table 1 (lung tissues).

#### Air liquid interface (ALI) cultures of HBECs

Human bronchial epithelial cells (HBECs) were obtained from endobronchial lining fluid (ELF) by minimally invasive bronchoscopic microsampling (BMS) from subsegmental airways for further investigation of indeterminate pulmonary nodules as described previously [Kahn et al. (2009)]. ELF was obtained from a noninvolved segment from the contralateral lung. Patients’ characteristics are shown in table 1 (ELF patients).

Primary HBECs were maintained in DMEM/F12 media (Gibco, Carlsbad, CA) supplemented with bovine pituitary extract (0.004 mL/mL), epidermal growth factor (10 ng/mL), insulin (5 μg/mL), hydrocortisone (0.5 μg/mL), triiodo-L-thyronine (6.7 ng/mL) and transferrin (10 μg/mL) (PromoCell), 30 nM sodium selenite (Sigma), 10 μM ethanolamine (Sigma), 10 μM phosphorylethanolamine (Sigma), 0.5 μM sodium pyruvate (Gibco), 0.18 mM adenine (Sigma), 15 mM Hepes (Gibco), 1x GlutaMAX (Gibco) and in the presence of 10 μM Rock-inhibitor (StemCell). After reaching confluency, cells were transferred to 12-well plates and plated at 90.000 cells/insert using the ThinCert™ with 0.4 μm pores (#665641, Greiner Bio-One). After 2-3 days, cells were airlifted by removing the culture from the apical chamber and PneumaCult-ALI media (StemCell) was added to the basal chamber only. Differentiation into a pseudostratified mucociliary epithelium was achieved after approximately 24-27 days. Afterwards, ALI cultures were characterized for their proper differentiation using Uteroglobin/CC10 (#RD181022220-01, BioVendor), Mucin 5AC (#ab3649, abcam), Keratin 5 (#HPA059479, Merck) and Tubulin beta 4 (#T7941, Merck). To do so, filters were fixed with 4% paraformaldehyde (PFA, Merck) and permeabilized with 0.1% Triton X-100 (Merck). First antibodies were applied over night at 4°C. Filters were incubated with Alexa Fluor 488 and 549 secondary antibodies (Dianova) for 40 minutes at 37°C. DNA was stained using Hoechst 33342 (#B2261, Merck). Pictures were taken using Leica SP5 confocal microscope and Leica Application Suite software (Leica). Pictures were assembled using Photoshop CS6 (Adobe).

### SINGLE NUCLEI AND SINGLE CELL RNA SEQUENCING

#### Single nuclei isolation (lung tissues)

Essentially, single nuclei were isolated as described elsewhere [Tosti et al. (2019)]. Briefly, snap frozen healthy lung tissue from lung adenocarcinoma patients was cut into pieces with less than 0.3 cm diameter and single nuclei were isolated at low pH by homogenizing the cells in 1 ml of citric acid-based buffer (Sucrose 0.25 M, Citric Acid 25 mM, Hoechst 33342 1 g/ml) at 4°C using a glass dounce tissue grinder. After one stroke with the “loose” pestle, the tissue was incubated for 5 minutes on ice, further homogenized with 3-5 strokes of the “loose” pestle, followed by another incubation at 4°C for 5 minutes. Nuclei were released by 5 additional strokes with the “tight” pestle and the nuclei solution was filtered through a 35 μm cell strainer. Cell debris was removed via centrifugation at 4°C for 5 minutes at 500x g, the supernatant was removed, followed by nuclei cell pellet resuspension in 700 μl citric-acid based buffer and centrifugation at 4°C for 5 minutes at 500x g. After carefully removing the supernatant, the nuclei cell pellet was resuspended in 100 μl cold resuspension buffer (25 mM KCl, 3 mM MgCl2, 50 mM Tris-buffer, 400 U RNaseIn, 1 mM DTT, 400 U SuperaseIn (AM2694, Thermo Fisher Scientific), 1g/ml Hoechst (H33342, Thermo Fisher Scientific). Nuclei concentration was determined using the Countess II FL Automated Cell Counter and optimal nuclei concentration was obtained by adding additional cold resuspension buffer, if needed. Subsequently, samples were processed using the 10x Chromium device with the 10x Genomics scRNA-Seq protocol v2 to generate cell and gel bead emulsions, followed by reverse transcription, cDNA amplification, and sequencing library preparation following the manufacturers’ instructions. Libraries have subsequently been sequenced one sample per lane on HiSeq4000 (Illumina; paired-end 26 x 74 bp)

#### Dissociation and Cryopreservation of ALI culture cells

Single cell suspensions were generated by adding 1x DPBS (Thermo, 14190094) to the apical chamber, followed by an incubation at 37°C for 15 minutes and subsequent washing of the cells with 1x DPBS (three times) to remove excess mucus. Adherent cells were dissociated via 0.25% Trypsin-EDTA (ThermoFisher Scientific, 25200056) treatment for 5 minutes at 37°C, followed by Dispase treatment (Corning, 354235) for 10 minutes at 37°C. Trypsinization was inactivated by adding cell culture medium with soybean trypsin inhibitor. Cells were spun down at 300x g for 5 minutes, resuspended in 1x DPBS, and passed through a 20 μm cell strainer (pluriSelect, 41-50000-03) to obtain a single cell solution and to remove cell clumps. Cells were spun at 300x g for 5 minutes and resuspended in cryopreservation medium (10% DMSO, 10% FBS, and 80% Culture Medium (PneumaCult-ALI) and frozen down gradually before shipping on dry-ice.

#### Single cell RNA sequencing library preparation (ALI cultures)

Cryopreserved cells were thawed at 37°C, spun down at 300xG for 5 minutes, the cell pellet was resuspended in 1x PBS with 0.05% BSA (Sigma-Aldrich, 05479), and passed through a 35 μm filter (Corning, 352235) to remove cell debris. Single cell suspensions were loaded onto the 10x Chromium device using the 10x Genomics Single Cell 3’ Library Kit v2 (10x Genomics; PN-120237, PN-120236, PN-120262) to generate cell and gel bead emulsions, followed by reverse transcription, cDNA amplification, and sequencing library preparation following the manufacturers’ instructions. The resulting libraries were sequenced with one sample per lane using the NextSeq500 (Illumina; high-output mode, paired-end 26 x 49 bp) or with two samples per lane using the HiSeq4000 (Illumina; paired-end 26 x 74 bp).

#### Pre-processing and data analysis

CellRanger software version 2.1.1 (10x Genomics) was used for processing of the raw sequencing data and the transcripts were aligned to the 10x reference human genome hg19 1.2.0. Low-quality cells were removed during pre-processing using Seurat version 3.0.0 (https://github.com/satijalab/seurat) based on the following criteria: (a) >200 or, depending on the sample, <6000 – 9000 genes (surgical lung tissues)/ <3000 – 5000 genes (ALI cultures), (b) <15% mitochondrial reads (Supp. Fig. 7). The remaining data were further processed using Seurat for lognormalization, scaling, merging, clustering, and gene expression analysis. Afterwards, all control samples were merged using the “FindIntegrationAnchors” and “IntegrateData” functions with their results being merged again and used for downstream analysis. Monocle3 [Cao et al. (2019)] was used infer cellular trajectories and dynamics. Dimensionality reduction, cell clustering, trajectory graph learning, and pseudotime measurement were performed with this tool.

### QUANTIFICATION AND STATISTICAL ANALYSIS

#### Statistical analyses

Statistical analyses were performed using R and Python 3.7.1 with scipy 0.14.1 and statsmodels 0.9.0. For comparisons between CPM values, the Wilcoxon rank sum test was used. To calculate the p-value for the overlap between sets, a hypergeometric test was employed. Expected molecule counts per cell were modelled using a binomial distribution and the observed detection probability. Gene set enrichment analysis were performed using Metascape Zhou et al. (2019)] on the KEGG, Canonical pathways, GO, Reactome, and Corum databases. P-values were adjusted for multiple testing using the Benjamini-Hochberg method Benjamini and Hochberg (1995)]. Boxes in box plots indicate the first and third quartile, with the median shown as horizontal lines. Whiskers extend to 1.5 times the inter-quartile-range. All individual data points are indicated on the plot.

## SUPPLEMENTAL INFORMATION

**Supp. Figure 1 – Exemplary characterization of an air liquid interface (ALI) culture derived from HBECs.** Filters of an ALI culture were stained for the indicated bronchial epithelial markers and analyzed using confocal microscopy.

**Supp. Figure 2 – Cell type identification by marker genes.** DotPlots indicate the percent expressing cells (size) and expression level (color) for selected marker genes for each cell type in HBECs.

**Supp. Figure 3 - Percentage of *ACE2*^+^/*TMPRSS2*^+^/*FURIt*^+^ positive cells**. Percentage of positive cells (defined here as having at least one read of the respective gene) for *ACE2, TMPRSS2,* and *FURIN* in primary lung and HBECs.

**Supp. Figure 4 – Quantification of ACE2+/TMPRSS2+ double positive cells.** Overlaps and enrichment statistics for all ACE2 and TMPRSS2 single and double positive cells in the primary lung and HBEC dataset. RF: representation factor, enrichment. P: hypergeometric tail probability.

**Supp. Figure 5 – Expression of DPP4 in primary lung.** Expression levels and percentage of positive cells for *DPP4* in the primary lung dataset.

**Supp. Figure 6 – Age-dependent cell type composition**. Cell type composition in the primary lung and HBEC dataset (age is color-coded).

**Supp. Figure 7 - single-cell sequencing quality control.** Numbers of unique molecular identifiers (UMIs, top row) and genes per cell are shown for each individual sample from primary lung tissue and HBECs.

## Notes

#### Summary of Updates

Corrected labels in supplementary figures; updated acknowledgments section.

https://doi.org/10.6084/m9.figshare.11981034.v1

https://data.mendeley.com/datasets/7r2cwbw44m/1

