## Supplementary figures and images for "SARS-CoV-2 receptor ACE2 and TMPRSS2 are predominantly expressed in a transient secretory cell type in subsegmental bronchial branches"

### Supplementary Figures

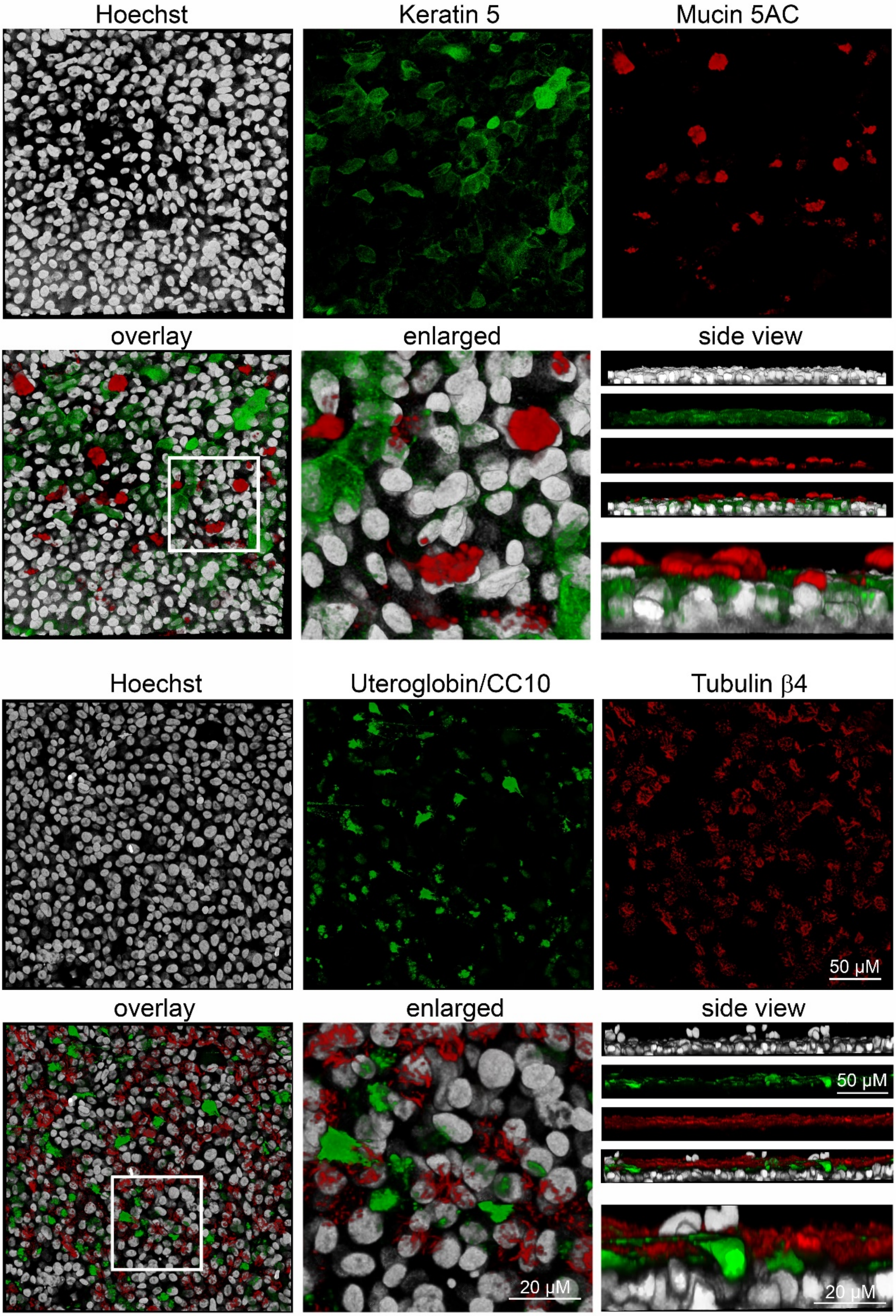

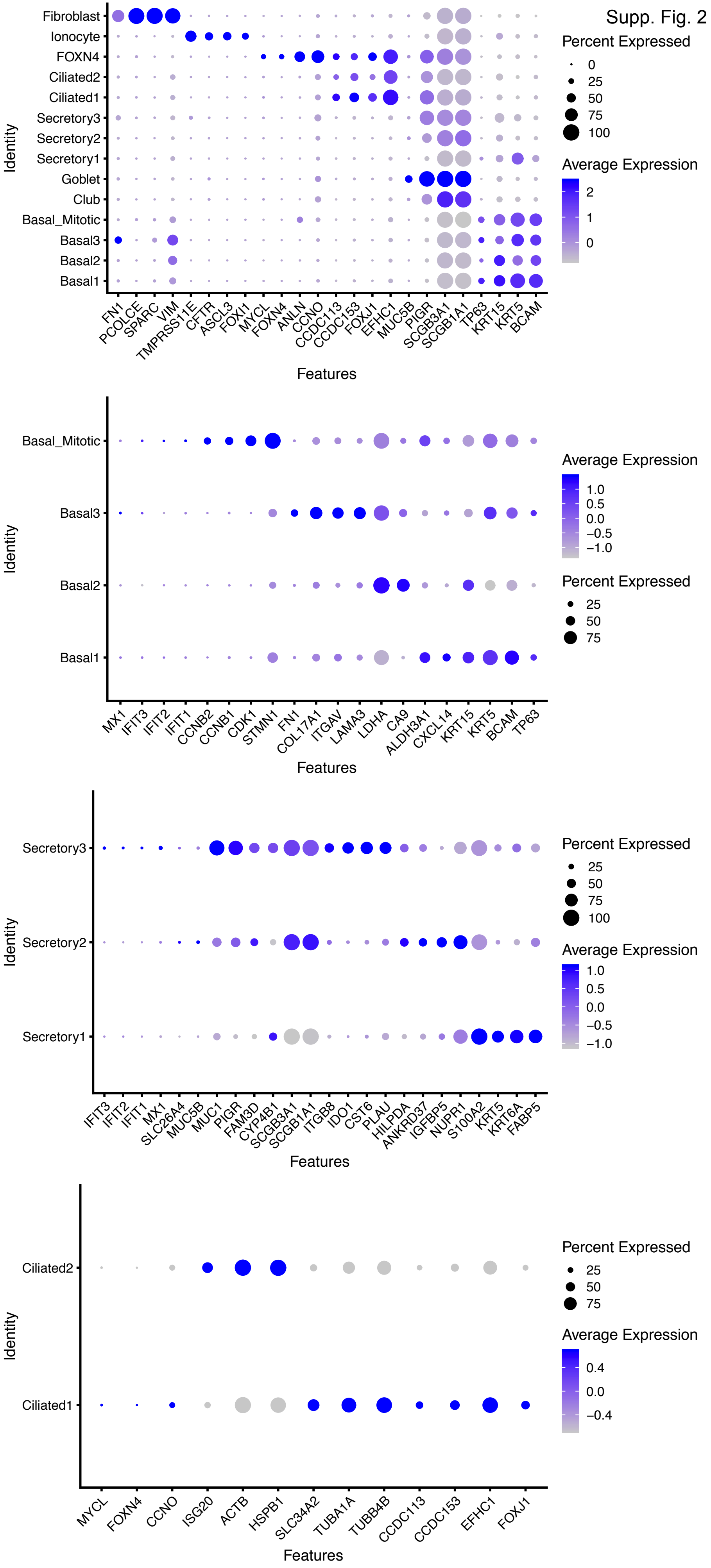

Supp. Fig. 3

lung HBECs

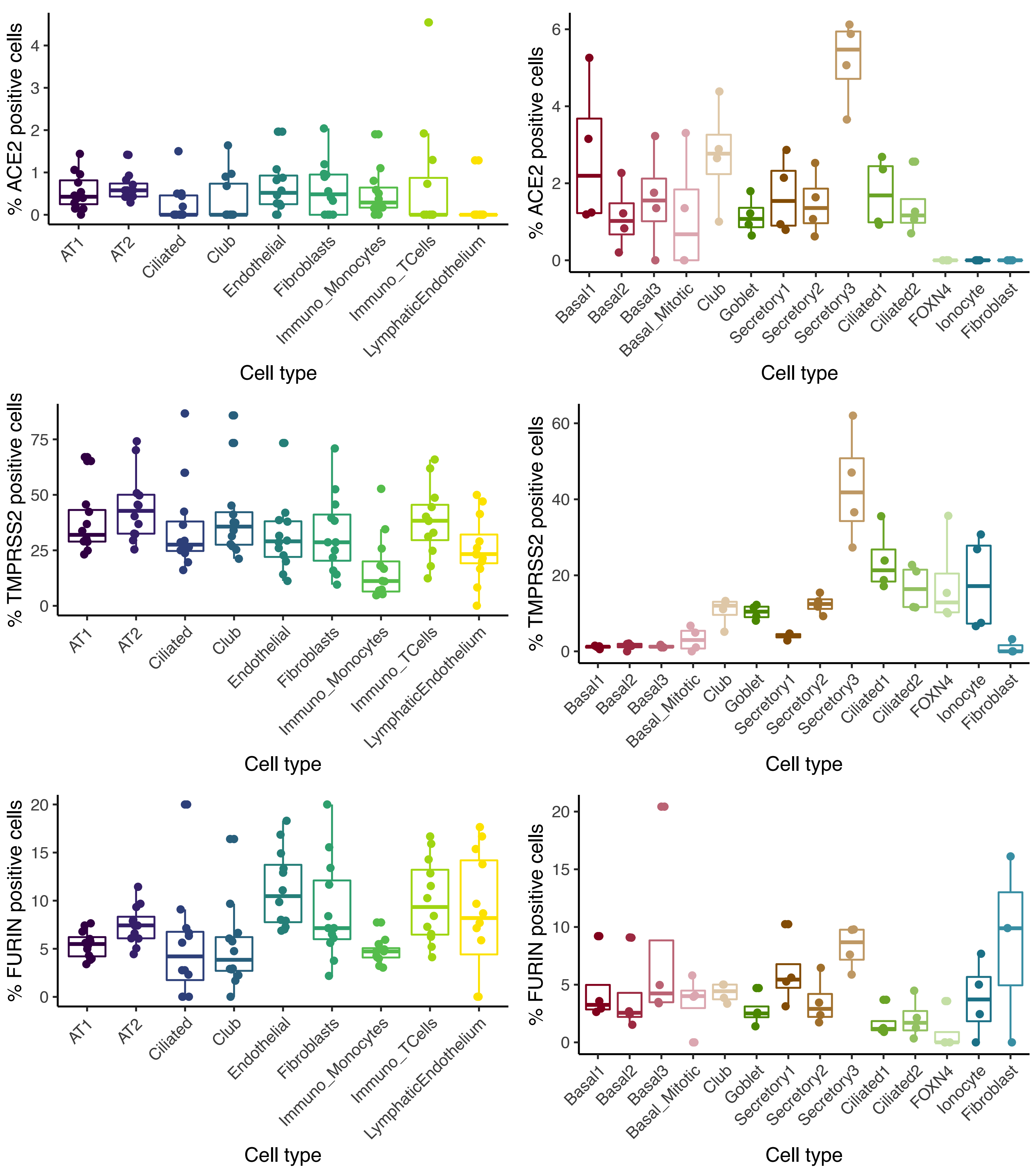

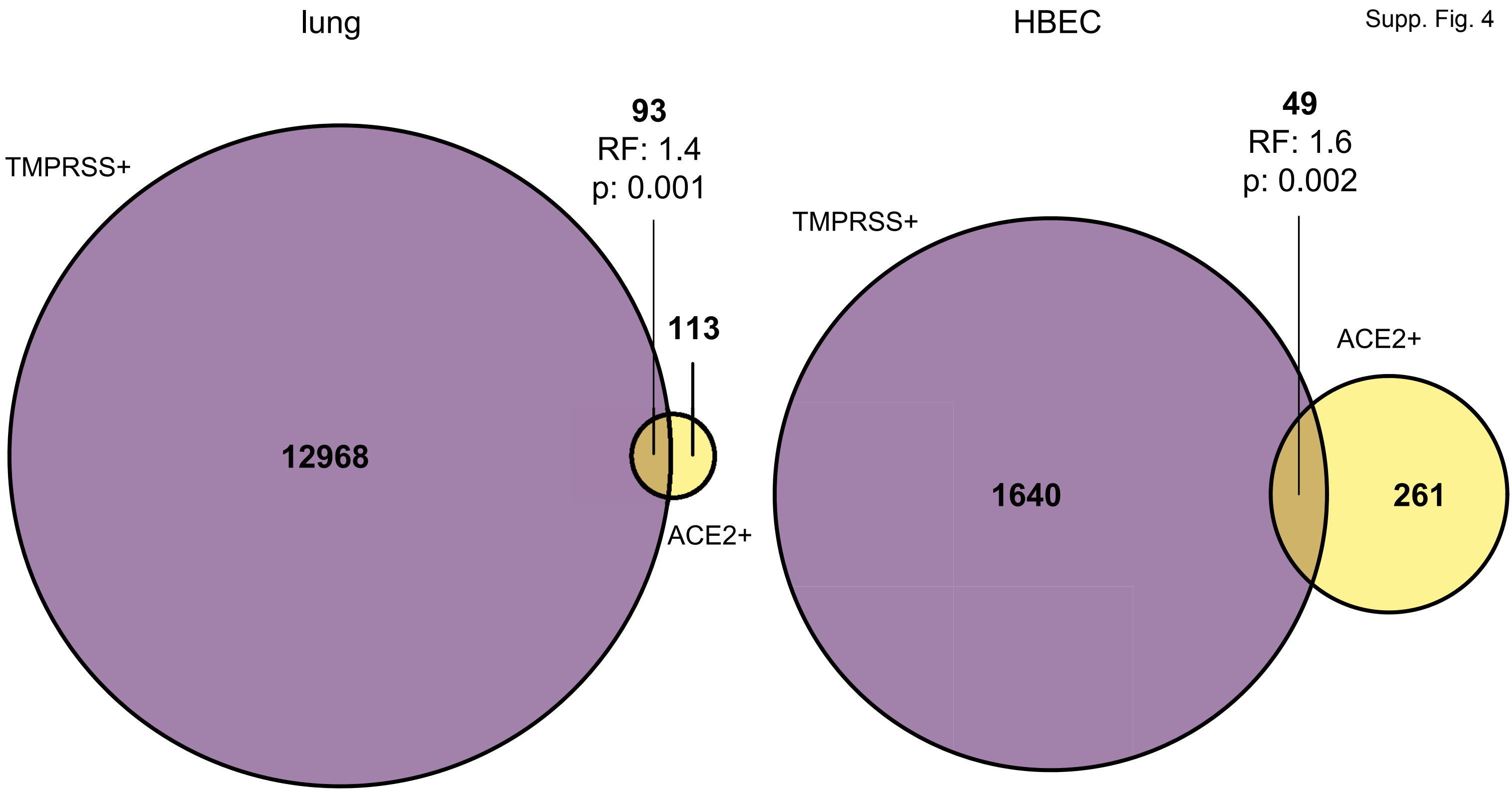

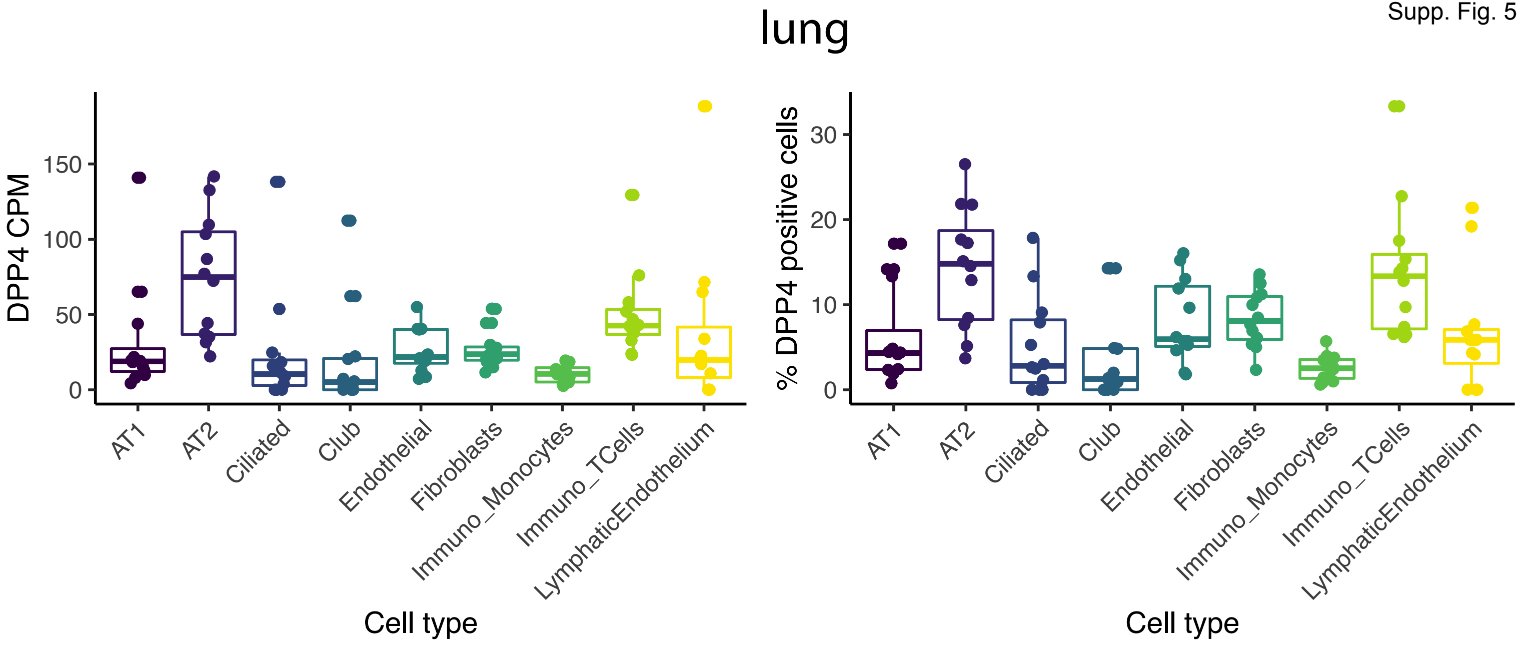

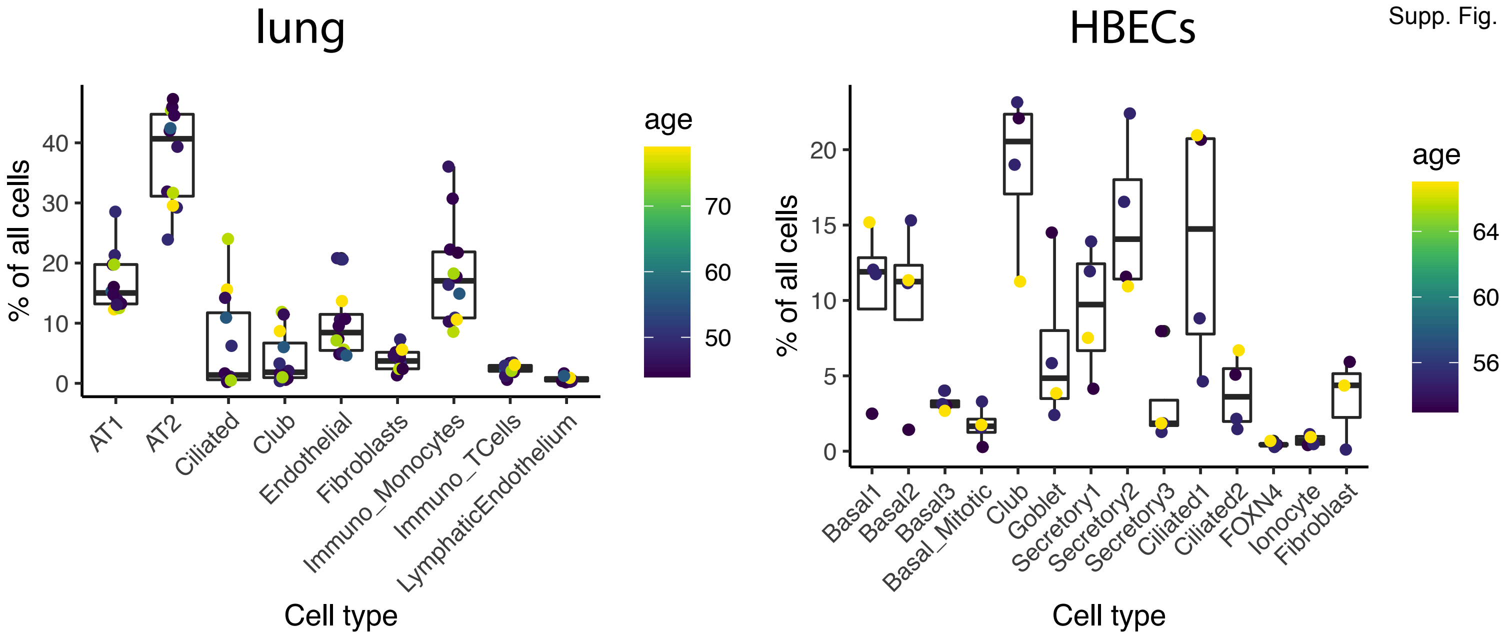

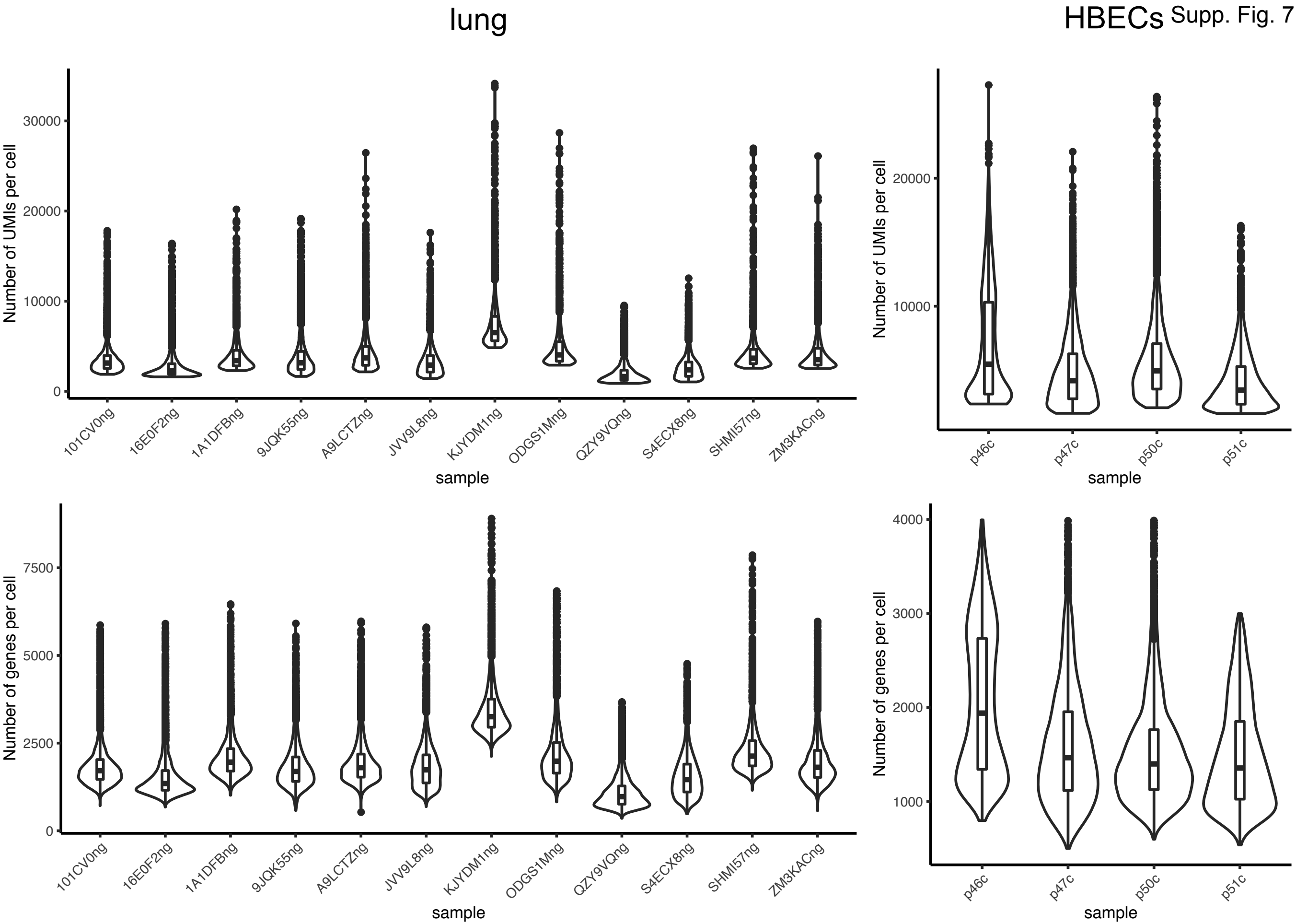
